# Activity budgets, social behavior, and fitness outcomes associated with a group fusion in baboons (*Papio cynocephalus* x *P. anubis*)

**DOI:** 10.64898/2026.04.30.721977

**Authors:** Brian A. Lerch, Maria J.A. Creighton, J. Kinyua Warutere, Jenny Tung, Elizabeth A. Archie, Susan C. Alberts

## Abstract

Many primates exhibit female philopatry and live in stable, female-bonded groups, where females are seldom exposed to novel same-sex conspecifics. Permanent group fusions are unique in bringing unfamiliar female conspecifics together but are rarely documented in these populations. We present a case study on a fusion of two groups from a hybrid population of baboons (*Papio cynocephalus* x *P. anubis*) living in the Amboseli basin of Kenya. We compared the behavior of eight adult females after the fusion to the behavior of females in groups that had not fused and compared pre- and post-fusion fitness outcomes. Following the fusion, the group’s activity budget and patterns of agonistic interactions were typical for the study population. Females preferred familiar grooming partners for a short period following the fusion; however, after three months, female grooming patterns were comparable to other groups, indicating rapid social integration. Females from the smaller pre-fusion group were lower ranking than females from the larger group. The fusion occurred following increased human-induced mortality in one of the two groups. From our limited sample, we observed no detectable fitness-related costs of fusion in terms of birth rates or offspring survival, while adult female mortality was low following the fusion. These results demonstrate the flexibility of female baboons in navigating exposure to novel same-sex conspecifics despite a species-typic pattern of female philopatry. Our results support previous research indicating that group fusions most likely occur when groups are too small to retain adult males, defend against predators, or compete with other groups.

## Introduction

Primates display a wide array of social structures and organizations that vary in dispersal patterns and the relative cohesiveness and stability of groups (Kappeler and van Schaik 2002; Koenig et al. 2013). A key dimension of this variation is the difference between species that live in groups that frequently undergo temporary fission and fusion events (i.e., “fission-fusion species”) versus those with low or non-existent fission-fusion dynamics (Aureli et al. 2008; Sueur and Maire 2014). Species with high frequencies of fissions and fusions routinely exhibit subgroup splitting and merging, requiring behavioral flexibility to manage shifting group compositions (Amici et al. 2008; Strier et al. 2014). In contrast, species with low frequencies of fissions and fusions maintain more stable groups, enabling consistency in social contact (Wrangham 1980; Port et al. 2020). In female-philopatric species where groups infrequently fuse, females are unlikely to encounter unfamiliar female groupmates as potential social partners at any point in their life, because all new females are born into the group (Henzi et al. 2000).

Much of what we know about fissions and fusions comes from species that exhibit frequent, temporary fissions and fusions. In these systems, fusions often lead to an increase in agonistic interactions, but such increases typically persist no longer than a few hours and are accompanied by a range of “greeting” behaviors that are thought to mitigate the social costs of fusing (Bauer 1974; Fedigan and Baxter 1984; Aureli and Schaffner 2007; Moscovice et al. 2015; Smith et al. 2015; Caselli et al. 2023). These patterns demonstrate that individuals living in fission-fusion systems are equipped with the behavioral flexibility to manage short-term changes in group composition. However, the reunion of groupmates in temporary fusions likely differs from permanent fusions of groups with unacquainted conspecifics. Unlike in species where temporary fissions and fusions are routine, group fusions in species with stable and discrete groups involve interactions between individuals without previous social relationships. Relatively little is known about how individuals respond to permanent fusions with novel conspecifics.

Many of the best studied cercopithecine primates live in relatively stable groups with low frequencies of fission-fusion events. Among these species, only approximately a dozen permanent group fusions have been documented, mostly in vervets (*Chlorocebus pygerythrus*), macaques (*Macaca* spp), and olive baboons (*Papio anubis*). Despite the rarity of fusion events in these species, common, though not universal, patterns have emerged. First, in most reported fusions, at least one of the groups involved in the fusion tends to be small with respect to the typical group size distribution for their species (Hauser et al. 1986; Isbell et al. 1991; Sugiura et al. 2002). Second, females from the smaller group tend to obtain the lowest ranks after the fusion (Dittus 1986; Hauser et al. 1986; Samuels et al. 1987; Isbell et al. 1991; Henzi et al. 2000; Sugiura et al. 2002; Jaffe and Isbell 2010; Strum 2012). Third, fusions often result when females follow adult males who emigrate and enter other groups, even though female dispersal is rare or otherwise absent in these species (Dittus 1986, 1987; Hauser et al. 1986; Isbell et al. 1991; Henzi et al. 2000; Strum 2012). Fourth, fusions often follow periods of increased mortality in one or both groups (Dittus 1987; Isbell et al. 1991; Takahata et al. 1994; Wasser et al. 2004; Jaffe and Isbell 2010; Strum 2012). These patterns of group fusion suggest that the causes of fusion are multifaceted: many factors must align to trigger a fusion, perhaps explaining their rarity in primates that otherwise live in stable and discrete groups (Dittus 1987). In general, permanent group fusions are thought to follow from conditions that favor large groups and render small groups unable to defend themselves, either from intergroup competition (Hauser et al. 1986; Isbell et al. 1991; Takahata et al. 1994) or predation (Jaffe and Isbell 2010; Strum 2012).

Because fusions are rare in primates that live in relatively stable groups with low fission-fusion dynamics, each additional documented fusion event meaningfully increases the total sample size available for understanding this uncommon behavior (Sugiura et al. 2002; Jaffe and Isbell 2010). Furthermore, theory suggests that fusions are demographically important: these events both reduce the total number of groups in a population and may also counter high mortality rates in small groups (Lerch and Abbott 2024). Lastly, by bringing unfamiliar females in contact, fusions provide insights into the behavioral flexibility of female primates who rarely need to establish social relationships with new, adult conspecifics. Based on the infrequency of permanent fusions reported in the literature, most females never experience these social conditions in their lifetimes.

Here, we report a fusion of two baboon groups living in the Amboseli ecosystem of Kenya. This population is admixed between yellow and olive (also called anubis) baboons (*Papio cynocephalus* x *P. anubis*), with majority yellow baboon ancestry (Tung et al. 2008; Vilgalys and Fogel et al. 2022). At the time of the fusion in 2012, the Amboseli baboon population had been under continuous observation for over 40 years. The extensive behavioral and demographic data on this study population provide an opportunity to compare behavior following the fusion to the population-level baseline, a comparison that, to our knowledge, has not been made previously for similar fusions. We asked three questions about the fusion, with the goal of understanding how otherwise philopatric females respond to living with novel same-sex conspecifics and quantifying the benefits that might explain why such permanent fusions occur. (1) How did female activity budgets and frequencies of agonistic interactions following the fusion compare to those of females in groups that did not experience recent fusions? (2) Was the fusion followed by an atypical distribution of social interactions among adult females? (3) Did females experience changes to demographic rates after the fusion? These questions allowed us to assess whether behavioral flexibility in the Amboseli baboons is sufficient to cope with uncommon, but large-scale changes in group composition. We focused our analyses on females because we do not routinely collect focal behavioral samples on adult males in this population (Alberts et al. 2024) and because, as the dispersing sex, males are accustomed to changing groups (Packer 1979; Alberts and Altmann 1995). Therefore, fusions are expected to present the most novel and unusual social challenges for females (Dittus 1986; Henzi et al. 2000). In addition, the fact that males regularly enter and exit the study population through dispersal makes it difficult to measure their survival and reproductive output.

## Methods

### The Fusion

The fusion of interest occurred in June of 2012 when two groups, Laza’s and Omo’s, fused to become one group, thereafter named Acacia’s group. At the time of the fusion, Omo’s group comprised 20 animals (five adult females, two adult males, and 13 dependent young and juveniles). Laza’s group was smaller than Omo’s group, with 14 animals (three adult females, three adult males, and eight dependent young and juveniles). Both groups were small compared to the average size of 62 animals for baboon groups in Amboseli (Southworth and Winans et al. 2026).

The fusion began when a prime-aged male, Buteo, left the smaller of the two groups, Laza’s, to join Omo’s group between June 19^th^ and June 23^rd^ of 2012. Buteo was followed by adult male Israel and adult female Luxin, the lowest-ranking female in Laza’s group, who were observed for the first time in Omo’s group on June 26^th^. The remaining members of Laza’s group included only one adult male, Alex, who was elderly and in poor condition. We first observed the rest of Laza’s group to have joined Buteo, Israel, and Luxin in Omo’s group on June 29^th^, signifying the establishment of the fusion product, “Acacia’s group” (Fig. 1). We considered the fusion complete on this day, given that all individuals from Laza’s group ceased ranging separately from Omo’s group from then on. This definition is consistent with several previous studies of group fusions (Dittus 1987; Jaffe and Isbell 2010), although Strum (2012) suggests a more stringent behavioral definition, which requires that grooming relationships become fully integrated before a fusion is considered complete. Following the fusion, Acacia’s group continued to use the former home range of Omo’s group (the larger of the two original groups), and females from the smaller group, Laza’s, became the lowest ranking females in Acacia’s group (Fig. 1).

**Fig. 1:**
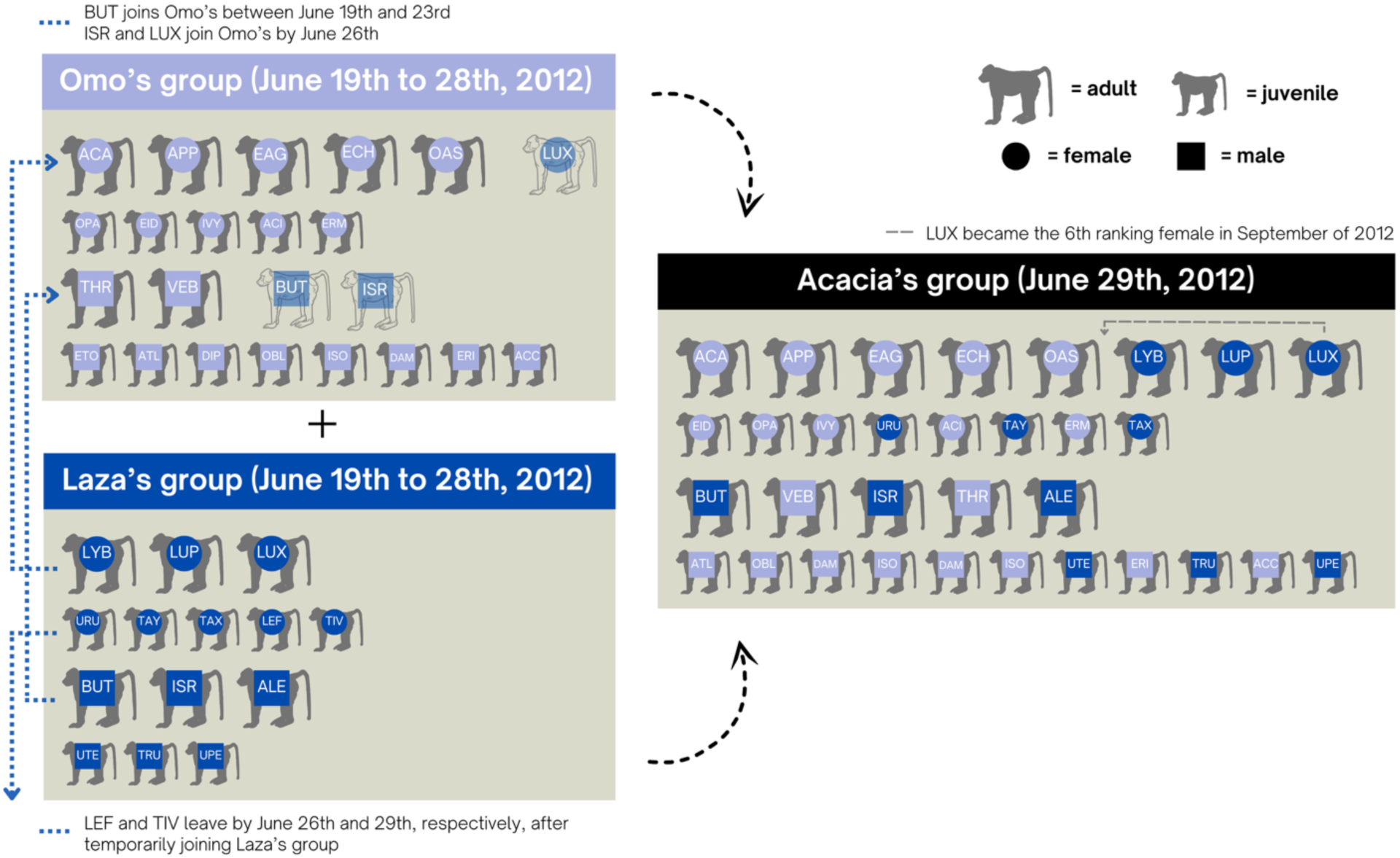
Demographics of two baboon groups in the Amboseli basin of Kenya before and after they fused in June of 2012. Color denotes origin group, shape denotes females versus males, and size denotes adults versus juveniles. Three letter codes refer to the abbreviation of individual names. Individuals are organized from left to right in order of dominance rank within each of the four age-sex classes (adult female, juvenile female, adult male, and juvenile male). Dominance ranks for the two pre-fusion groups, Omo’s and Laza’s, come from the last month that rank information was available for each group prior to being dropped as study groups (May of 2011 and December of 2011, respectively). In four cases, individuals were missing rank information before the fusion because they either weren’t born yet (ERM) or weren’t present in the group (VEB, LEF, TIV) in the month when ranks were last recorded. These individuals are not in rank order and are listed last in their respective rows.

### Study Subjects

Our subjects were wild baboons studied by the Amboseli Baboon Research Project (ABRP) in the Amboseli ecosystem of Kenya (Alberts and Altmann, 2012). Experienced ABRP observers collect behavioral and demographic data year-round on a near-daily basis, and all study animals are individually known through visual recognition (Alberts et al. 2024). The subjects of interest in this study were the adult females living in Laza’s and Omo’s groups when the 2012 fusion occurred.

Omo’s group originated in April of 1999 as the result of a fission of a larger group and existed as a stable group (i.e., all demographic changes stemmed from births, deaths, or immigration or emigration of individual males) for 11 years prior to the fusion. In contrast, Laza’s group existed for only eight months prior to the fusion, after originating from the fission of a larger group in October of 2011 (Southworth and Winans et al. 2026). The matrilines in Laza’s and Omo’s groups had been separated since before the onset of continuous data collection in 1971.

### Data Collection

ABRP observers visit study groups on a near-daily basis to record which members are present on that day, as well as any births and deaths. We have not observed permanent emigration for infants or for adult or juvenile females in this population; therefore, we infer deaths once an infant or female is missing for multiple days and is not located in the surrounding area (Alberts et al. 2024). We track adult female reproductive states using visual cues including the presence of sexual swellings or menstrual bleeding, and changes in the paracallosal skin (which become pink in color during pregnancy) (Alberts et al. 2024).

For groups under intensive study, observers also conduct focal sampling of individual behavior. During ten-minute focal samples, observers follow known juveniles or adult females in a pre-determined randomized order and record their activity (e.g., resting, walking, feeding, or grooming) at one-minute intervals known as point samples. These focal samples can be used to calculate individual activity budgets. While completing group censuses and conducting focal sampling of specific individuals, observers additionally collect data on all grooming and agonistic interactions that occur in their line of sight (Alberts et al. 2024). We used these data to compare the relative frequency and directionality of post-fusion grooming and agonistic interactions within and between members of the two original groups to other groups studied by the ABRP. Agonistic interactions were also used to determine dominance ranks before and after the fusion. In female baboons, dominance ranks are determined primarily through matrilineal rank inheritance (i.e., females nepotistically inherit social ranks from their mothers), but females sometimes deviate from this expectation (Lee and Oliver 1979; Silk 2009; Lea et al. 2014).

Demographic data were available for both Omo’s group and Laza’s group prior to the fusion, as censuses of Omo’s group occurred one to four times per month while censuses of Laza’s group occurred approximately three times per week. However, behavioral data during this time-period were sparse, because the ABRP dropped Omo’s group as a study group in May of 2011 and was in the process of dropping Laza’s group when the fusion occurred. Therefore, the focal samples important for estimating activity budgets and collecting grooming and agonistic interactions were limited or absent (Fig. S1). The ABRP last recorded dominance ranks for Omo’s and Laza’s groups in May of 2011 and December of 2011, respectively, 13 and six months before the fusion. Because of these behavioral data gaps, we could not directly compare behavioral patterns before and during the fusion. Near-daily observations resumed on the newly formed Acacia’s group after the fusion, with the resumption of regular demographic and behavioral data collection on June 29^th^ of 2012 (Fig. S1) and focal data collection one month later, on August 1^st^ of 2012. Even with these data limitations, this fusion is the only example from the Amboseli baboons for which systematically collected data exist.

### How did female activity budgets and frequencies of agonistic interactions following the fusion compare to those of females in groups that did not experience recent fusions?

To assess whether female activity was typical following the fusion, we compared the activity budgets and the relative frequencies of agonistic interactions for females in Acacia’s group in August, September, and October of 2012 (following the fusion in June of 2012) to those of females in other study groups (using data from 1999 onwards). To control for potential seasonal effects, we restricted our data sets for the other study groups to the same three months of the year, which represent the end of the long dry season in Amboseli and can have marked effects on ecology and behavior (Alberts et al. 2005).

We first used focal samples to calculate the percentage of total time females in any given group spent resting, walking, feeding, and grooming each August, September, and October of any given year and plotted these values against group size. We then evaluated if Acacia’s group showed typical patterns for its size (i.e., if its data point in August, September, and October fell within the same range as other groups). We followed a similar procedure to assess whether the relative per capita frequencies of agonistic interactions were typical following the fusion. However, in this case, we calculated the relative frequency of agonistic behaviors rather than the proportion of focal time dedicated to them. We used this approach for agonistic interactions because their short duration means they are not recorded as behavioral states but instead as events during focal samples, and also *ad lib* throughout daily observations.

We could not assess activity budgets or frequencies of agonistic interactions for Acacia’s group in June or July of 2012 (directly after the fusion) because the focal data collection necessary for calculating activity budgets and controlling for observer effort had not yet begun on this group. Therefore, our comparisons began in August of 2012, approximately five weeks after the fusion was completed. Consequently, we could not assess changes in female activity budgets during the first five weeks after the fusion and, as a result, we may have missed transient increases in rates of aggression, as observed in two previous studies of group fusion in cercopithecine primates (Dittus 1987; Jaffe and Isbell 2010).

### Was the fusion followed by an atypical distribution of social interactions among adult females?

Even if the fusion did not appreciably change activity budgets or relative rates of agonistic interactions, it could have affected who interacted with whom. For example, females could preferentially groom their original groupmates compared to those from the other group or primarily direct aggression towards females from the other group compared to their original groupmates. To investigate these possibilities, we assessed the number of grooming and agonistic interactions between females who originated in different pre-fusion groups versus the same pre-fusion group in the three months following the fusion (June 29^th^ to September 29^th^ of 2012; even though focal sampling did not begin until August, we had more detailed data on dyadic social interactions collected during censuses prior to this date). We then compared these values to the expected values if interactions were distributed among females irrespective of their pre-fusion origin.

Because all females in Omo’s group became dominant to all Laza’s females, any patterns of directionality in agonistic and grooming interactions could simply reflect typical patterns of social interactions among high-ranking and low-ranking females. Therefore, we also tested whether the observed patterns in grooming and agonistic interactions deviated from typical differences between low- and high-ranking females, in support of these differences being unique to the post-fusion environment. To do so, we first identified seven other times over the history of the ABRP where a group had the same number of adult females as Acacia’s group did between June 29^th^ and September 29^th^ of a given year. For each of these groups, we then assessed the frequency of interactions that occurred within and between the five highest-ranking and the three lowest-ranking females within the three-month window, mirroring the rank divide between Omo’s and Laza’s females after the fusion.

To test if the distribution of agonistic interactions between females in Acacia’s group differed from other groups after accounting for typical rank effects, we calculated the total number of female-female agonistic interactions in Acacia’s group and each of the seven other groups with the same number of females. We then performed four linear regressions, where the number of total female-female agonistic interactions recorded in each of the eight groups predicted the number of female-female agonistic interactions in which: (i) one of the five highest-ranking females won interactions with one of the three lowest-ranking females, (ii) one of the three lowest-ranking females won interactions with one of the five highest-ranking females, (iii) one of the five highest-ranking females won interactions with one of the five highest-ranking females, and (iv) one of the three lowest-ranking females won interactions with one of the three lowest-ranking females. Our choice to split females into groups of three and five was based on the split represented in the fusion, where the three lowest ranking adult females in Acacia’s group (the fusion group) were from Laza’s group and the five highest ranking adult females were from Omo’s. We also used this approach to evaluate whether grooming patterns among females in Acacia’s group post-fusion were unusual after accounting for typical rank effects. If the data points for Acacia’s group fell far from the regression lines, this would suggest that the post-fusion environment had an atypical distribution of interactions among females for a group of its size, even after accounting for rank-based differences.

### Did females experience changes to demographic rates after the fusion?

To assess whether females experienced detectable changes in fitness associated with the fusion we compared per capita birth rate, infant survival to one year, and adult female survival before and after the fusion. To do so, we compared annual outcomes for these metrics in Omo’s and Laza’s groups before the fusion to Acacia’s group after the fusion. Laza’s group only existed as an independent group for less than one year before the fusion while Omo’s became its own group in 1999; thus, all but one data point from before the fusion came from Omo’s group.

We measured birth and adult death rates for each year as the number of events per adult female day and the infant death rates for each year as the number of infant deaths per infant day, where an infant or female day represented a day where a given female or infant was considered a member of the group of interest (note that this approach still results in a single estimate per group, per year). We measured annual birth and death rates as the rate of events per day, as opposed to events per year, because births, infant aging, sexual maturation, and deaths changed the infant and adult female composition of a group, and thus the denominator for rate calculations, by the day. Calculating events per female or infant day allowed us to account for this daily demographic variation, which in turn affected the probability of observing a birth or death. Because birth and death rates are low for baboons and we only had data from one fusion, we did not have enough power to test for statistical differences in demographic rates before and after the fusion. Therefore, we plotted the rate of births and deaths per day for each year and calculated an average of these rates across years before the fusion (from 1999 through the pre-fusion part of 2012, including both Omo’s data points for all years and Laza’s data points for the single year in which that group existed). We then compared these rates to those in Acacia’s group after the fusion (from the post-fusion part of 2012 through 2023) to identify any qualitative evidence of shifts in births and deaths in relation to the fusion.

In addition to direct measures of these fitness proxies, we examined whether Acacia’s group was an outlier for its group size in the number of recorded injuries and pathologies observed per individual in the three months following the fusion. Injuries (including wounds, visible limps, and broken bones) and pathologies (including both chronic and acute illnesses) are documented by field observers as part of the standard ABRP observation protocol (Alberts et al. 2024). These records can reveal periods of increased fighting (in the case of injuries) and increased disease transmission (in the case of sickness cues) and thus serve as an indicator of individual health.

### Data Availability Statement

Data and code have been made available through the Duke Research Data Repository and can be accessed at https://doi.org/10.7924/r4r561.

## Results

### How did female activity budgets and frequencies of agonistic interactions following the fusion compare to those of females in groups that did not experience recent fusions?

The activity budgets of females in Acacia’s group following the fusion were similar to other groups in the population during the same three months of the year, considering all years in the study. Following the fusion, neither the percentage of time Acacia’s females spent on common behaviors (grooming, feeding, resting, and walking) nor the relative frequency of per capita agonistic interactions were atypical, suggesting that fusion-associated changes in behavioral patterns were absent during August, September, or October (Fig. 2). While the relative frequency of agonistic interactions fell in the upper quartile of the population for all three months (Fig. 2a), such a pattern occurred in 7.4% of group-years from August through October when considering all groups in our dataset. One month, September, was also an outlier for multiple activities: during the September after the fusion, females in Acacia’s group spent less time grooming (in the bottom 9.8% of group-months; Fig. 2b), more time feeding (in the top 2.5% of group-months; Fig. 2c), and less time resting (in the bottom 2.2% of group-months; Fig. 2d) relative to other groups. These patterns may suggest higher foraging demand in the post-fusion month of September 2012 than is typical for groups of the same size. However, August was not an outlier for these same behaviors, even though it was closer to the fusion (Fig. 2b, 2c, and 2d). Thus, in the absence of further information (e.g., data on food availability), it is difficult to draw conclusions about the cause of the fused group’s unusual activity budget in September.

**Fig. 2:**
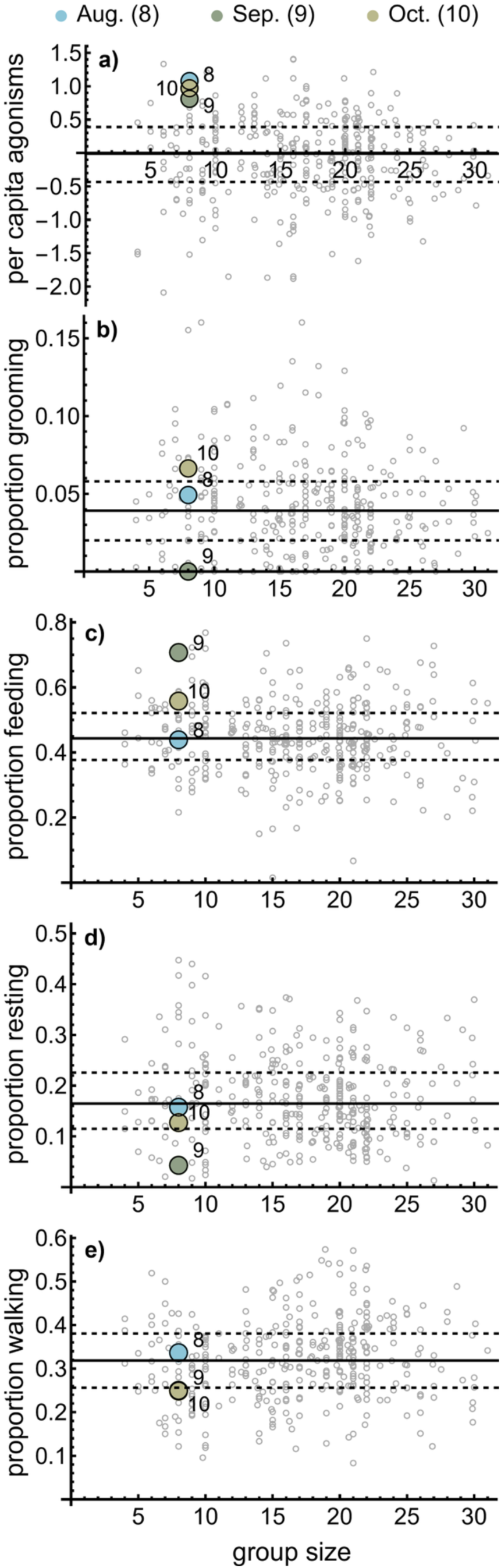
Time spent on different behaviors by adult female baboons in the Amboseli basin of Kenya each August, September, and October from 1999 to 2023 as a function of group size. (a) Relative frequency of per capita agonistic interactions between adult females, controlled for observer effort (the number of focal samples divided by the adult female group size). Specifically, the y-axis is the residual of a regression of the number of agonistic interactions per capita versus observer effort. Only interactions involving aggression are included (i.e., displacements and submission-only interactions are excluded. The following panels show the activity budget as the proportion of time spent (b) grooming, (c) feeding, (d) resting, and (e) walking. Colored points represent Acacia’s group in the three months after it formed via fusion once focal sampling resumed: 8=August, 9=September, and 10=October. Note that in panel d) the data points for September and October fall completely on top of each other. Horizontal lines show the median value (solid) and the interquartile range (dashed).

### Was the fusion followed by an atypical distribution of social interactions among adult females?

Both grooming and agonistic interactions among Omo’s and Laza’s females were distributed non-randomly in the three months immediately after the fusion (*χ*-squared test: *χ*^2^ = 14.2, df = 3, *p* = 0.003 and *χ*^2^ = 164.1, df = 3, p < 0.001, respectively; Table 1). Females from Laza’s group (the smaller of the two pre-fusion groups) lost more decided agonistic interactions with Omo’s females than expected by chance (Fig. S2). Laza’s females also groomed Omo’s females less often than expected by chance (Table 1, Fig. S2). Luxin, the first of Laza’s females to join Omo’s group, was responsible for the majority (5 of 7) of total grooming events directed from Laza’s females to Omo’s highest-ranking female, Acacia (the namesake for the post-fusion group; Fig. S3). This is notable because Luxin ascended in rank relative to her former groupmates: she went from being the lowest ranking female in Laza’s to become the sixth ranking female in Acacia’s group three months after the fusion (Fig. 1), placing her higher in rank than all other females originating from Laza’s group.

**Table 1:** *χ*-squared test indicates that social interactions were distributed non-randomly between June 29^th^ and September 29^th^ of 2012 after the fusion of Laza’s and Omo’s groups, with the null expectation that interactions between females from each group would be proportional to the number of dyads with a given origin; i.e., because there were 5 females in Omo’s group, 3 in Laza’s group, and individuals cannot interact with themselves, the expected proportions were: Omo-to-Omo: 20/56, Omo-to-Laza: 15/56, Laza-to-Omo: 15/56, and Laza-to-Laza: 6/56. Interaction types indicate either i) a female who originated in the first listed group groomed a female who originated in the second listed group, or ii) a female who originated in the first group won an agonistic interaction with a female who originated in the second group.

| Interaction type | Omo-to-Omo count | Omo-to-Laza count | Laza-to-Omo count | Laza-to-Laza count | p-value |
| --- | --- | --- | --- | --- | --- |
| Grooming | 56 | 28 | 15 | 12 | 0.003 |
| Agonism | 61 | 133 | 1 | 15 | <0.001 |

Despite the apparent imbalance in the direction of interactions following the fusion, we found that after accounting for differences in social ranks, the distribution of agonistic interactions among high- and low-ranking females in Acacia’s group directly after the fusion were consistent with from those in other groups. In other words, Acacia’s group fell on the trendline of expected interaction rates when taking into account the overall number of recorded agonistic interactions per group (Fig. 3, all panels). This result suggests that the high frequency of submissions observed from Laza’s females (reported in Table 1) can be explained by typical rank-related differences in the direction of agonistic interactions, as opposed to being unique to the post-fusion environment.

**Fig. 3:**
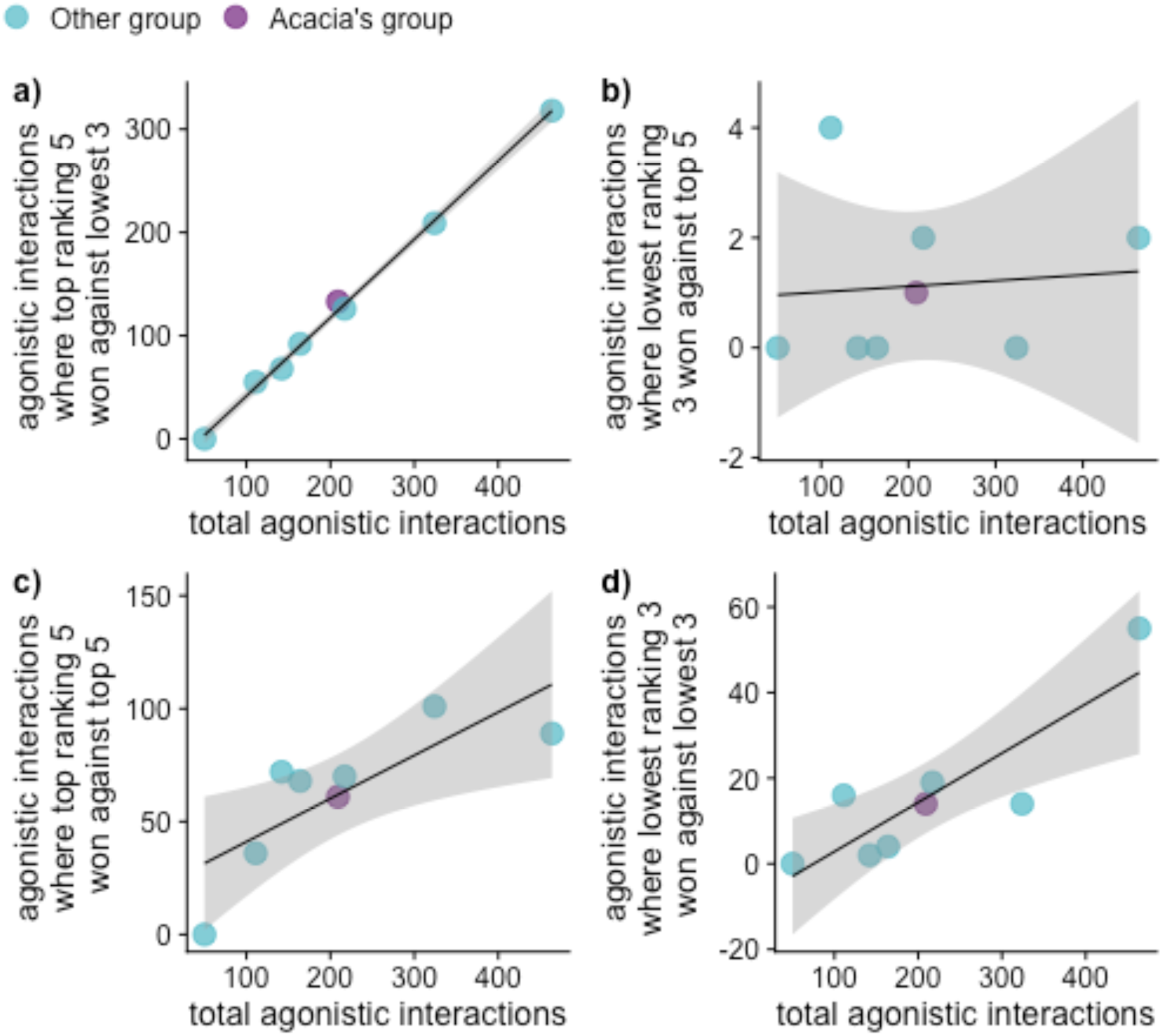
Relationship between total agonistic interactions and agonistic interactions between specific subsets of group members, based on rank, within groups containing eight adult female baboons between June 29^th^ and September 29^th^ of a given year (between 1972 and 2023) in the Amboseli basin of Kenya: a) one of the five highest-ranking females won interactions with one of the three lowest-ranking females, b) one of the three lowest-ranking females won interactions with one of the five highest-ranking females, c) one of the five highest-ranking females won interactions with one of the five highest-ranking females, and d) one of the three lowest-ranking females won interactions with one of the three lowest-ranking females. Acacia’s group in the three months after it formed via fusion is plotted in purple while other groups are plotted in blue. The solid black line shows the linear regression trendline from a regression where the number of agonistic interactions falling within a particular category was regressed on the total number agonistic interactions. The shaded gray area shows the 95% confidence interval around the regression line. Comparisons are made between the top five and bottom three females because the three lowest ranking adult females in Acacia’s group were in Laza’s group prior to the fusion and the five highest ranking adult females were in Omo’s group prior to the fusion (Fig. 1). Note that the vertical axis scale changes across panels.

Unlike patterns in agonistic interactions, grooming patterns among females in Acacia’s group differed noticeably from groups that had not undergone a fusion, after accounting for rank differences. Females from Laza’s group (the lowest three ranking) groomed females from Omo’s group (the highest five ranking) less frequently than low-ranking females groom high-ranking females in other groups containing the same number of females (Fig. 4b). Specifically, the data point for Acacia’s group was 2.2 standard deviations below the predicted value given by the regression line (Fig. 4b). Meanwhile, females from Omo’s group (the highest five ranking) groomed each other more frequently than high ranking females from groups of the same size, such that the data point for Acacia’s group was 2.0 standard deviations above the predicted value given by the regression line (Fig. 4b). These biases in grooming were absent by four to six months following the fusion (Fig. S4), suggesting that these atypical patterns were specifically related to the time-period directly following the fusion. These results suggest that while patterns of rank-related aggression between the two subsets of females were not unusual following the fusion, social integration between females from the two pre-fusion groups was not achieved until a few months following the fusion.

**Fig. 4:**
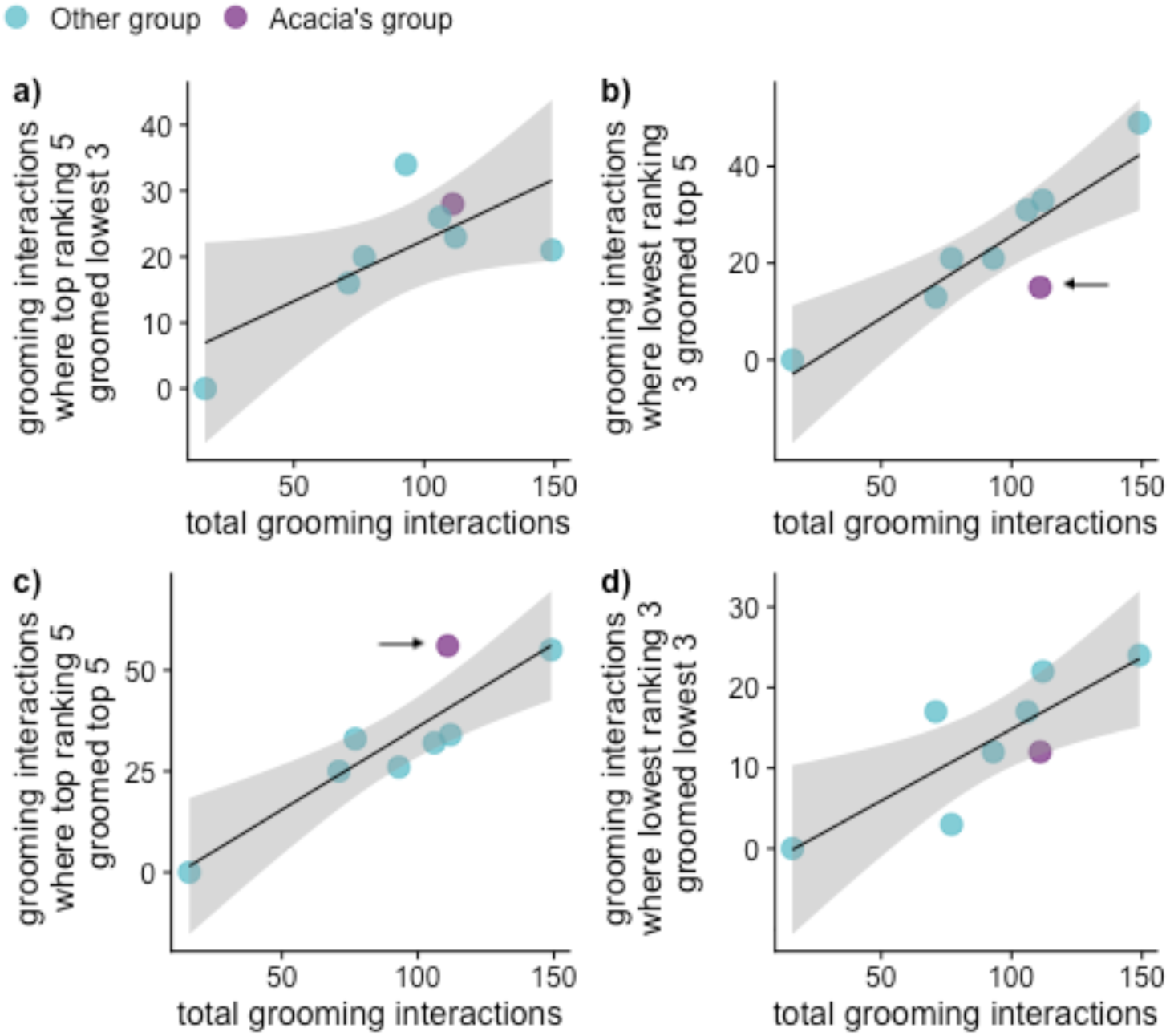
Relationship between total grooming interactions and grooming interactions between specific subsets of group members, based on rank, in groups containing eight adult female baboons between June 29^th^ and September 29^th^ of a given year (between 1972 and 2023) in the Amboseli basin of Kenya: a) one of the five highest-ranking females groomed one of the three lowest-ranking females, b) one of the three lowest-ranking females groomed one of the five highest-ranking females, c) one of the five highest-ranking females groomed one of the five highest-ranking females, and d) one of the three lowest-ranking females groomed one of the three lowest-ranking females. Acacia’s group in the three months after it formed via fusion is plotted in purple while other groups are plotted in blue. The solid black line shows the linear regression trendline from a regression where the number of grooming interactions falling within a particular category was regressed on the total number grooming interactions. The shaded gray area shows the 95% confidence interval around the regression line. Comparisons are made between the top five and bottom three females because the three lowest ranking adult females in Acacia’s group were in Laza’s group prior to the fusion and the five highest ranking adult females were in Omo’s group prior to the fusion (Fig. 1). Arrows point to cases where Acacia’s group in the months following the fusion deviated meaningfully from the regression line. Note that the vertical axis scale changes across panels.

### Did females experience changes to demographic rates after the fusion?

There was no evidence of increased injuries or pathologies following the fusion, contrary to expectations if the fusion led to health costs by inciting physical conflict or disease transmission. No injuries or pathologies were recorded for adult females in Acacia’s group during the three months following the fusion (Fig. 5), and the frequency of injuries and pathologies for all group members (including males and juveniles) were consistent with groups of a similar size (Fig. S5).

**Fig. 5:**
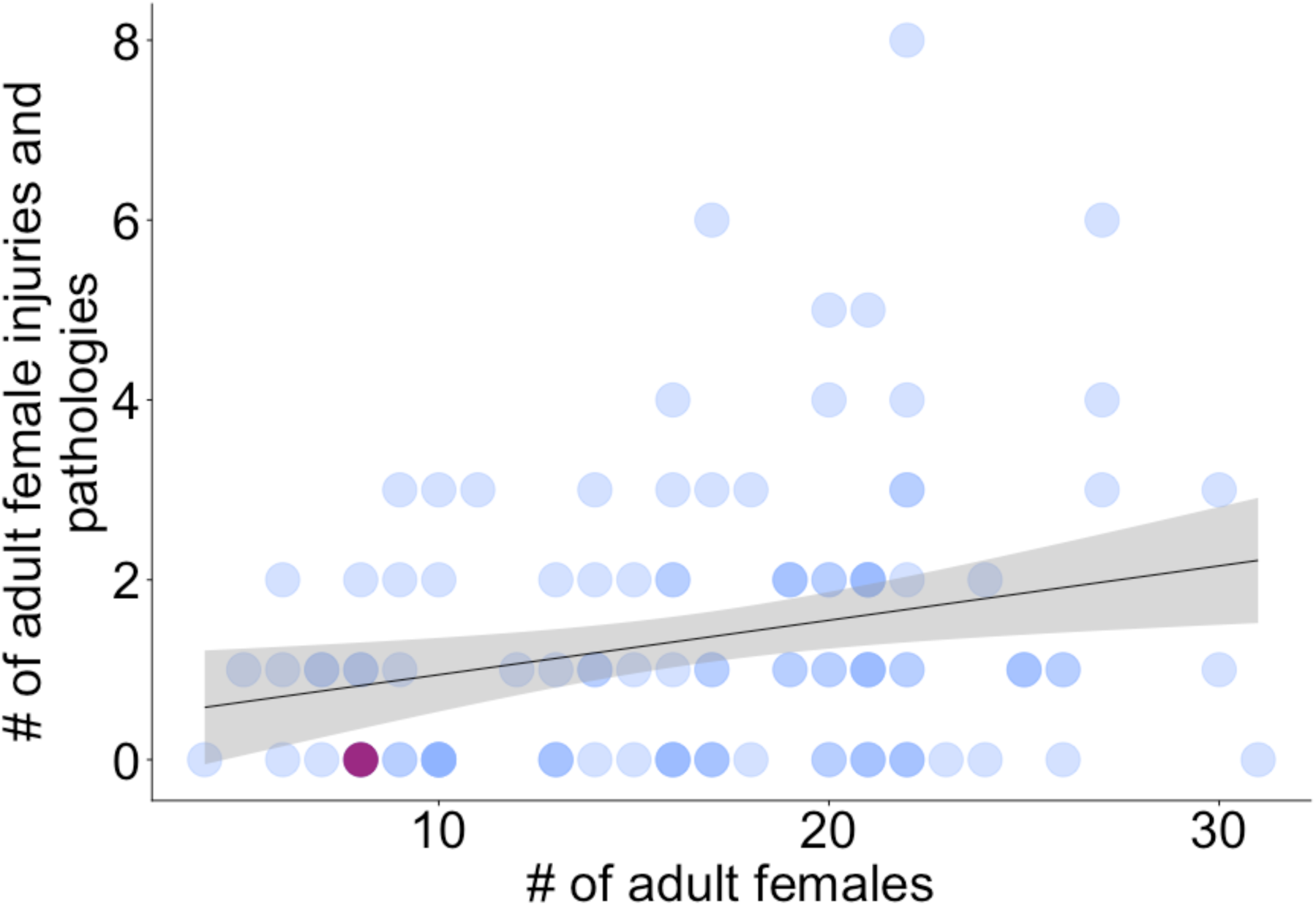
Number of recorded injuries and pathologies affecting adult female baboons in a group between June 29^th^ and September 29^th^ of a given year (from 2000 to 2023) in the Amboseli basin of Kenya versus the total number of adult females in the group. Acacia’s group in the three months after it formed via fusion is plotted in purple while other group-year combinations are plotted in blue.

Birth rate (the number of infants born per adult female per day) did not notably change after the fusion, suggesting that the fusion carried neither costs nor benefits to reproductive output. The mean annual birth rate was 1.21 x 10^-3^ births per adult female day before the fusion compared to 1.42 x 10^-3^ births per adult female day after the fusion, corresponding to a negligible increase from 0.442 to 0.519 births per adult female per year (Fig. 6a). Infant mortality was slightly higher after the fusion, suggesting that the fusion did not improve offspring survival. The mean annual infant death rate (the number of infant deaths per infant day) was 6.03 x 10^-4^ deaths per infant day before the fusion (each infant had a 19.8% chance of dying in its first year) and 9.04 x 10^-4^ deaths per infant day after (each infant had a 28.1% chance of dying in its first year) (Fig. 6b). Infant mortality appeared to be particularly high in Acacia’s group the year of the fusion. However, this apparent spike in mortality was driven by one infant death coupled with the low birth rates immediately before the fusion (Fig. 6b) and we thus interpret it with caution.

**Fig. 6:**
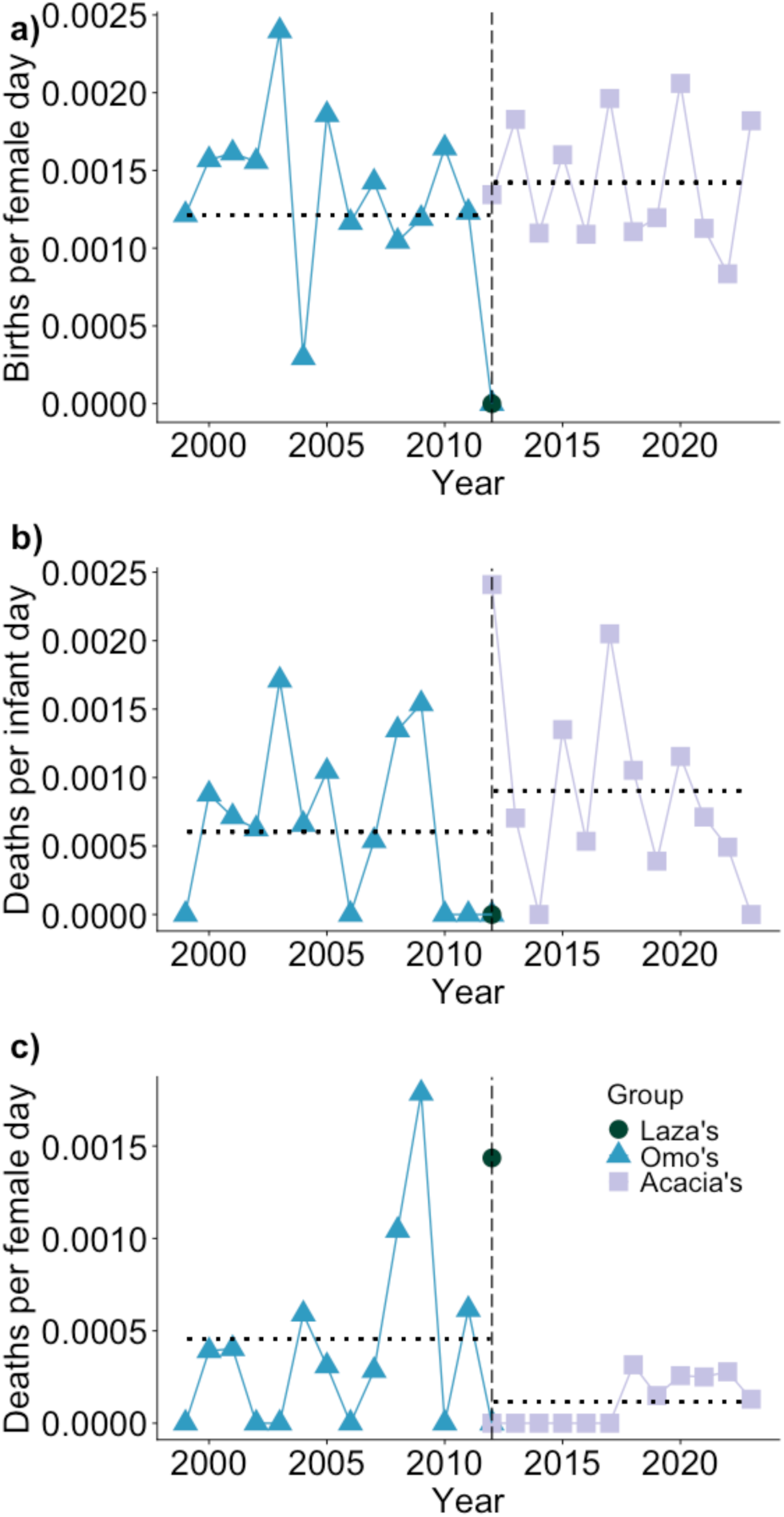
Rate of births and deaths per day during each calendar year from 1999 to 2023, for two baboon groups in the Amboseli basin of Kenya involved in a fusion and their fusion product, Acacia’s group. a) Birth rate: number of infants born per adult female per day. b) Infant death rate: number of infant deaths per infant day. c) Adult female death rate: number of female deaths per female day. A female day represents a day where a female was considered a member of the group of interest. An infant day represents a day where an infant was considered a member of the group of interest. The vertical dashed line shows the year the groups fused. Horizontal dotted lines show means before and after the fusion. Note that Laza’s group only has one data point as they fissioned from a larger group in 2011.

Consistent with the possibility that the fusion enhanced fitness by promoting adult survival, the adult female death rate (the number of female deaths per female day) in Omo’s and Laza’s groups before the fusion was nearly four times higher than the death rate after the fusion in Acacia’s group (4.57 x 10^-4^ deaths per female day compared to 1.15 x 10^-4^ deaths per female day). This change in death rate corresponded to a 15.4% adult female yearly mortality rate before the fusion and a 4.1% adult female yearly mortality rate after the fusion. Part of this change can be attributed to a large spike in mortality in Omo’s group in 2009, three years prior to the fusion, when three adult females died in a span of four days due to conflict with humans (Fig. 6c). Two adult males were also killed in this same event. In addition to the events in 2009, most years before the fusion exhibited a non-zero number of adult female deaths, and the pre-fusion death rate is still more than three times higher than the post-fusion death rate, even if 2009 was excluded from the calculation. In Laza’s group, one of four adult females died in the one year during which the group existed before the fusion (i.e., after it was formed through the fission of another group in 2011), resulting in a high female death rate before the fusion (Fig. 6c). The large spike in death rate for Laza’s group caused by this single female death reflects the sensitivity of the y-axis to demographic stochasticity in small groups, including both Laza’s and Omo’s. Nonetheless, it is noteworthy that no adult female deaths occurred in Acacia’s for six years following the fusion (Fig. 6c).

## Discussion

Analyzing data from a rare case of group fusion in the Amboseli baboons, we show that two formerly distinct groups can socially integrate relatively rapidly with minimal disruptions to activity budgets or adult female social interactions. This rapid integration is possible even though stable groups with low fission-fusion dynamics are the norm in this population and adult female baboons typically have little experience in navigating exposure to novel same-sex conspecifics. In particular, one month after the fusion there were no detectable effects on female activity budgets or the relative frequency or patterns of agonistic interactions. A transient bias towards grooming previous groupmates was evident in the three months after the fusion, a pattern reported in other fusions (Hauser et al. 1986; Suguira et al. 2002; Jaffe and Isbell 2010), but this bias disappeared after three months. Subsequently, the low dominance ranks of the females that originated in Laza’s group (the smaller of the two groups) was the only behavioral trace of the two original groups. While other studies have also observed rapid integration after group fusion (Dittus 1987; Jaffe and Isbell 2010), this pattern is not universal: full behavioral integration of a fused group took more than two years to complete in a population of olive baboons (Strum 2012).

The demographics of Acacia’s fusion closely paralleled other documented examples of group fusions in primates with a similar social organization. The fusion included at least one small group (as in Hauser et al. 1986; Isbell et al. 1991; Sugiura et al. 2002); in this case both groups were small compared to the population mean. Furthermore, evidence suggests that the fusion occurred when adult female and juvenile group members from the smaller group followed adult males who immigrated into the larger group (as in Dittus 1986, 1987; Hauser et al. 1986; Isbell et al. 1991; Henzi et al. 2000; Strum 2012). Two of three males in Laza’s group joined Omo’s group days before other group members, with only an elderly male in poor physical condition remaining in Laza’s group. Adult males serve several important functions in baboon groups including mating, protection from predators, and protection of females and their young from new immigrant males (Cowlishaw 1994; Weingrill 2000; Nguyen et al. 2009; Moscovice et al. 2015). Thus, the loss of young or prime adult males likely represents a serious risk to females.

Also consistent with previous studies, females from the smaller of the two groups, Laza’s, became the lowest ranking individuals in Acacia’s group (as in Dittus 1986; Hauser et al. 1986; Samuels et al. 1987; Isbell et al. 1991; Henzi et al. 2000; Sugiura et al. 2002; Jaffe and Isbell 2010; Strum 2012). Because female baboons exhibit nepotistic rank inheritance, the low ranks that females from a smaller group assume after a baboon group fusion will be transferred to their daughters. Furthermore, as higher-ranking matrilines grow, the ordinal rank position of these low-ranking females can become even lower (as observed in spotted hyenas, *Crocuta crocuta*, Strauss and Holekamp 2019). For instance, Luxin, who ranked 6 of 8 females soon after the fusion, ranked 16 of 21 in December 2023, eleven and a half years after the fusion; in other words, a larger absolute number of females outranked her in 2023 than did immediately following the fusion. The pattern in which the ‘joining females’ become low ranking may be related to patterns observed in captive studies of rhesus macaques (*Macaca mulatta*), in which the order of introduction into an enclosure determines female rank order (Jarrell et al. 2008; Tung et al. 2012; Snyder-Mackler et al. 2016). Specifically, females in the smaller of two parent groups who accept the rank superiority of females in the larger group may be following a behavioral rule of thumb similar to that seen in captivity, where females accept the rank superiority of females already present in an enclosure. In captive settings, the first females introduced into an enclosure may have more opportunities to establish social alliances and thus maintain a higher rank (Jarrell et al. 2008). These patterns are consistent with Luxin’s rise in ranks following her early dispersal into Omo’s group and social affiliation (i.e., grooming) with the dominant female, Acacia. Indeed, coalitionary support — from either adult females or adult males — may be broadly important for rank reversals in a range of animal species (Strauss and Holekamp 2019) and is known to reinforce female rank in our study population (Silk et al. 2004). Unfortunately, the sparsity of coalitionary data in the period following the fusion meant we could not investigate the role of coalitions in determining rank outcomes in this study.

Previously documented fusions in cercopithecine primates with similar patterns of social organization have also reported elevated mortality risk preceding fusion events (Dittus 1987; Isbell et al. 1991; Takahata et al. 1994; Wasser et al. 2004; Jaffe and Isbell 2010; Strum 2012). Our limited data here are consistent with these observations, in support of the idea that primates in small groups may pursue group fusions as a strategy to reduce mortality by increasing group size. Increased predation risk is one of the main threats to small groups (Stacey 1986; Hill and Lee 1998; Bettridge and Dunbar 2013) and has been argued to trigger group fusion (Wasser et al. 2004; Jaffe and Isbell 2010; Strum 2012). Because human-induced mortality is likely experienced as predation by the baboons, the fusion of Omo’s and Laza’s groups relatively soon after the human-induced mortality spike in Omo’s group in 2009 is consistent with this argument. In some primate species, groups with relatively few adult males — which are typically small groups — experience a high risk of infanticide from immigrating males (Crockett and Janson 2000; see Zipple et al. 2017 for an analysis of infanticide in Amboseli). We speculate that the combined increase in the risk of both predation and infanticide could serve as a self-reinforcing mechanism that keeps small groups small and can lead to gradual decline in membership and increased risk of group extinction. Fusion presents a way to bypass the suppressed growth of small groups and prevent this outcome (Lerch and Abbott 2024).

One other fusion occurred in Amboseli in 1972 during a brief lapse in observations (McCuskey 1975; Altmann 1980; Samuels et al. 1987) and before the current behavioral data collection protocols were implemented, making information on that fusion very limited. However, qualitative information about that fusion also matches patterns from Acacia’s group and other documented cercopithecine fusions. This older fusion occurred in September of 1972 when Hightail’s group (with approximately nine members plus infants) and Alto’s group (with approximately 31 members plus infants) fused into one group. An obvious similarity between the 1972 fusion, the 2012 fusion, and fusions in other studies of related primates is the involvement of a relatively small group. In the year leading up to the 1972 fusion, the smaller group (Hightail’s) also experienced a very high death rate: in August of 1971 it contained 14 members, and by August of 1972 it contained only nine members and had experienced seven deaths in that 12-month period. This death rate is nearly double that of the larger groups in the study population at the time (McCuskey 1975). At least one of these deaths, involving adult female Corrie, was caused by predation. Thus, increased mortality may have been a motivating factor in this fusion. Another similarity between the 1972 and 2012 fusions is that adult females from Hightail’s group (the smaller group in 1972) obtained the lowest rank positions in the adult female hierarchy after the fusion (Samuels et al. 1987).

In more than fifty years of study, including 24 baboon groups in Amboseli, only two group fusions have occurred, underscoring the rarity of these events in this population. As argued by others, the rarity of group fusions in species that typically exhibit a low frequency of fission-fusion dynamics make each additional report valuable for building a generalizable understanding of the conditions that lead to fusions, and the responses of, and consequences for, the individuals who experience them (Dittus 1986; Henzi et al. 2000; Sugiura et al. 2002; Jaffe and Isbell 2010). The rarity of group fusions also suggests that several conditions must align to trigger these events (Dittus 1987). The combination of factors that appear important in the decision to fuse — small group size, male emigration, and high mortality — further suggests that most instances of group fusion that have been observed, including the two reported here in the Amboseli baboons, are a strategy motivated by risk avoidance. Risk avoidance could also explain why ‘joining females’ in a group fusion are willing to accept low dominance ranks in their new groups. However, even the general conditions that seem to predict fusions in baboons and their close relatives are probably not universal. In contrast to the hypothesis that group fusions are motivated by risk avoidance, one group fusion in olive baboons may be better explained by improved access to food resources (Strum 2012). Specifically, when a smaller group joined a larger one, the post-fusion group eventually gained access to an area with richer resources. Both infant and adult mortality increased in the year following the fusion, but in subsequent years both survival and reproductive rates improved. More data from populations that experience a surplus in resources (e.g., in human-modified environments) would help resolve the relative importance of between-group tolerance versus reduction in risk in motivating primate group fusions.

The behavioral flexibility required for baboons to undergo group fusion in a relatively rapid and peaceful manner underscores the argument that species that exhibit low versus high fission-fusion dynamics lie on a continuum (Aureli et al. 2008) and, as social and demographic conditions dictate, even primates who typically inhabit stable groups can move along this continuum with ease. Given the apparent ability for primates to handle changes to their social organization, it is not surprising that dramatic changes in species-typical patterns of social organization have evolved relatively quickly in primates. For example, the baboon genus is only ∼2 million years old but includes species with extreme male reproductive skew, species with strong and enduring male coalitions and relatively low reproductive skew, and species with multilevel societies (Fischer et al. 2019). This wide range of social patterns seen between species, and the great behavioral flexibility within species, exemplified by our case study on group fusion in the Amboseli baboons, sheds light on the success of the primate order over macroevolutionary timescales. In the most extreme case, this flexibility may also help account for the remarkable ability of humans to have rapidly and successfully spread across the globe (Richerson and Boyd 2006; Boyd et al. 2011; Henrich 2015).

## Acknowledgements

We gratefully acknowledge the support of the National Science Foundation and the National Institutes of Health, through multiple grants over the years, including currently active NIH grants R01AG071684, R01AG075914, and R61AG078470. We also thank the Max Planck Institute for Evolutionary Anthropology, Duke University, Princeton University, and the University of Notre Dame for financial and logistical support. M.J.A.C. was supported by the Natural Sciences and Engineering Research Council of Canada (PGSD3 – 577867 – 2023). In Kenya, our research was approved by the Wildlife Research Training Institute (WRTI), Kenya Wildlife Service (KWS), the National Commission for Science, Technology, and Innovation (NACOSTI), and the National Environment Management Authority (NEMA). We additionally thank the University of Nairobi, the Kenya Institute of Primate Research (KIPRE), the National Museums of Kenya, the members of the Amboseli-Longido pastoralist communities, the Enduimet Wildlife Management Area, Ker & Downey Safaris, Air Kenya, and Safarilink for their cooperation and assistance in the field. Particular thanks go to Jeanne Altmann for her foundational contributions to the Amboseli Baboon Research Project dataset, to the Amboseli Baboon Research Project long-term field team (R.S. Mututua, S. Sayialel, J.K. Warutere, I.L. Siodi, and L. Musembei), and to Nairobi-based support team T. Wango and V. Oudu. Thanks also to Kerri Smith for her data collection on Acacia’s group following the fusion. The baboon project database, Babase, is expertly managed by J. Gordon and C. Broderick. Database design and programming are provided by K. Pinc. This research was approved by the IACUCs at Duke University and University of Notre Dame and the Ethics Council of the Max Planck Society, and adhered to all the laws and guidelines of Kenya. For a complete set of acknowledgments of funding sources, logistical assistance, and data collection and management, please visit http://amboselibaboons.nd.edu/acknowledgements/.

## Notes

https://doi.org/10.7924/r4r561

### Competing Interest Statement

The authors have declared no competing interest.

### Summary of Updates

This updated version reflects changes to the manuscript title and structure, along with minor changes to some analyses.

## References

Alberts, S. C., and J. Altmann. 1995. Balancing costs and opportunities: dispersal in male baboons. The American Naturalist 145:279–306.

Alberts, S. C., and J. Altmann. 2012. The Amboseli Baboon Research Project: 40 years of continuity and change. In P. M. Kappeler & D. P. Watts (Eds.), Long-term field studies of primates (pp. 261–287). Springer, Berlin, Heidelberg.

Alberts, S. C., E. A. Archie, J. Altmann, and J. Tung. 2024. Monitoring guide for the Amboseli Baboon Research Project. See https://amboselibaboons.nd.edu/downloads/.

Alberts S.C., J. Hollister-Smith, R. S. Mututua, S. N. Sayialel, P. M. Muruthi, J. K. Warutere, J. Altmann. 2005. Seasonality and long term change in a savannah environment. In: D. K. Brockman, C. P. van Schaik (Eds), Primate Seasonality: Implications for Human Evolution (pp. 157–196). Cambridge University Press.

Altmann, J. 1980. Baboon Mothers and Infants. Harvard University Press, Cambridge, MA.

Amici, F., F. Aureli, and J. Call. 2008. Fission-fusion dynamics, behavioral flexibility, and inhibitory control in primates. Current Biology 18:1415–1419.

Aureli, F., and C. M. Schaffner. 2007. Aggression and conflict management at fusion in spider monkeys. Biology Letters 3:147–149.

Aureli, F., C. M. Schaffner, C. Boesch, S. K. Bearder, J. Call, C. A. Chapman, … and C. P. van Schaik. 2008. Fission-fusion dynamics: New research frameworks. Current Anthropology 49:627–654.

Bauer, H. R. 1974. Behavioral changes about the time of reunion in parties of chimpanzees in the Gombe Stream National Park. Proceedings of the International Congress of Primatology 5:295–303.

Bettridge, C. M., and R. I. M. Dunbar. 2013. Predation as a determinant of minimum group size in baboons. Folia Primatologica, 83:332–352.

Boyd, R., P. J. Richerson, and J. Henrich. 2011. The cultural niche: Why social learning is essential for human adaptation. Proceedings of the National Academy of Sciences 108:10918– 10925.

Caselli, M., B. Malaman, G. Cordoni, J.-P. Guéry, J. Kok, E. Demuru, and I. Norscia. 2023. Not lost in translation: Changes in social dynamics in Bonobos after colony relocation and fusion with another group. Applied Animal Behaviour Science 261:105905.

Cowlishaw, G. 1994. Vulnerability to predation in baboon populations. Behaviour 131:293–304.

Crockett, C. M., and C. H. Janson. 2000. Infanticide in red howlers: female group size, male membership, and a possible link to folivory. In C. P. van Schaik & C. H. Janson (Eds.), Infanticide by males and its implications (pp. 75–98). Cambridge University Press, Cambridge, UK.

Dittus, W. P. J. 1986. Sex differences in fitness following a group take-over among toque macaques: Testing models of social evolution. Behavioral Ecology and Sociobiology 19:257–266.

Dittus, W. P. J. 1987. Group fusion among wild toque macaques: An extreme case of inter-group resource competition. Behaviour 100:247–291.

Fedigan, L. M., and M. J. Baxter. 1984. Sex differences and social organization in free-ranging spider monkeys (*Ateles geoffroyi*). Primates 25:279–294.

Fischer, J., J. P. Higham, S. C. Alberts, L. Barrett, J. C. Beehner, T. J. Bergman, … and D. Zinner. 2019. The natural history of model organisms: Insights into the evolution of social systems and species from baboon studies. eLife 8:e50989.

Hauser, M. D., D. L. Cheney, and R. M. Seyfarth. 1986. Group extinction and fusion in free-ranging vervet monkeys. American Journal of Primatology 11:63–77.

Henrich, J. 2015. The secret of our success: How culture is driving human evolution, domesticating our species, and making us smarter. Princeton University Press, Princeton, NJ, USA.

Henzi, S. P., L. Barrett, A. Weingrill, P. Dixon, and R. A. Hill. 2000. Ruths amid the alien corn: Males and the translocation of female Chacma baboons. South African Journal of Science 96:61–62.

Hill, R. A., and P. C. Lee. 1998. Predation risk as an influence on group size in cercopithecoid primates: Implications for social structure. Journal of Zoology 245:447–456.

Isbell, L. A., D. L. Cheney, and R. M. Seyfarth. 1991. Group fusions and minimum group sizes in vervet monkeys (*Cercopithecus aethiops*). American Journal of Primatology 25:57–65.

Jaffe, K. E., and L. A. Isbell. 2010. Changes in ranging and agonistic behavior of vervet monkeys (*Cercopithecus aethiops*) after predator-induced group fusion. American Journal of Primatology 72:634–644.

Jarrell, H., J. B. Hoffman, J. R. Kaplan, S. Berga, B. Kinkead, and M. E. Wilson. 2008. Polymorphisms in the serotonin reuptake transporter gene modify the consequences of social status on metabolic health in female rhesus monkeys. Physiology & Behavior 93:807–819.

Kappeler, P. M., and C. P. van Schaik. 2002. Evolution of primate social systems. International Journal of Primatology 23:707–740.

Koenig, A., C. J. Scarry, B. C. Wheeler, and C. Borries. 2013. Variation in grouping patterns, mating systems and social structure: what socio-ecological models attempt to explain. Proceedings of the Royal Society B 268:20120348.

Lea, A. J., N. H. Learn, M. J. Theus, J. Altmann, and S. C. Alberts. 2014. Complex sources of variance in female dominance rank in a nepotistic society. Animal Behaviour 94:87–99.

Lee, P. C., and J. I. Oliver. 1979. Competition, dominance and the acquisition of rank in juvenile yellow baboons (*Papio cynocephalus*). Animal Behaviour 27:576 –585.

Lerch, B. A., and K. C. Abbott. 2024. A flexible theory for the dynamics of social population: Within-group density dependence and between-group processes. Ecological Monographs 94:e1604.

McCuskey, S. A. 1975. Demography and behavior of one male groups of yellow baboons (*Papio cynocephalus*). University of Virginia, M.S. Thesis.

Moscovice, L. R., T. Deschner, and G. Hohmann. 2015. Welcome back: Responses of female bonobos (*Pan paniscus*) to fusions. PLOS One 10:e0127305.

Nguyen, N., R. C. Van Horn, S. C. Alberts, and J. Altmann. 2009. “Friendships” between new mothers and adult males: Adaptive benefits and determinants in wild baboons (*Papio cynocephalus*). Behavioral Ecology and Sociobiology 63:1331–1344.

Packer, C. 1979. Inter-troop transfer and inbreeding avoidance in *Papio anubis*. Animal Behaviour 27:1–36.

Port, M., H. Hildenbrandt, I. Pen, O. Schülke, J. Ostner, and F. J. Weissing. 2020. The evolution of social philopatry in female primates. American Journal of Physical Anthropology 173:397–410.

Richerson, P. J., and R. Boyd. 2006. Not by genes alone: How culture transformed human evolution. University of Chicago Press, Chicago, IL, USA.

Samuels, A., J. B. Silk, and J. Altmann. 1987. Continuity and change in dominance relations among female baboons. Animal Behaviour 35:785–793.

Silk, J. B. 2009. Nepotistic cooperation in non-human primate groups. Philosophical Transactions of the Royal Society B: Biological Sciences 364:3243–3254.

Silk, J. B., S. C. Alberts, and J. Altmann. 2004. Patterns of coalition formation by adult female baboons in Amboseli, Kenya. Animal Behaviour 67:573–582.

Smith, J. E., J. R. Estrada, H. R. Richards, S. E. Dawes, K. Mitsos, and K. E. Holekamp. 2015. Collective movements, leadership and consensus costs at reunions in spotted hyaenas. Animal Behaviour 105:187–200.

Snyder-Mackler, N., J. N. Kohn, L. B. Barreiro, Z. P. Johnson, M. E. Wilson, and J. Tung. 2016. Social status drives social relationships in groups of unrelated female rhesus macaques. Animal Behaviour 111:307–317.

Southworth, C. A., J. C. Winans, J. B. Gordon, N. H. Learn, W. A. Wilber, C. Andreadis, … and J. Tung. 2026. Demographic, behavioral, and ecological data from a long-term field study of wild baboons in Amboseli, Kenya. Scientific Data 13:311.

Stacey, P. B. 1986. Group size and foraging efficiency in yellow baboons. Behavioral Ecology and Sociobiology 18:175–187.

Strauss, E. D., and K. E. Holekamp. 2019. Social alliances improve rank and fitness in convention-based societies. Proceedings of the National Academy of Sciences 116:8919–8924.

Strier, K. B., P. C. Lee, and A. R. Ives. 2014. Behavioral flexibility and the evolution of primate social states. PLOS One 9:e114099.

Strum, S. C. 2012. Darwin’s monkey: Why baboons can’t become human. Yearbook of Physical Anthropology 55:3–23.

Sueur, C., and A. Maire. 2014. Modelling animal group fission using social network dynamics. PLOS One 9:e97813.

Sugiura, H., N. Agetsuma, and S. Suzuki. 2002. Troop extinction and female fusion in wild Japanese macaques in Yakushima. International Journal of Primatology 23:69–84.

Takahata, Y., S. Suzuki, N. Okayasu, and D. Hill. 1994. Troop extinction and fusion in wild Japanese macaques of Yakushima Island, Japan. American Journal of Primatology 33:317– 322.

Tung, J., L. B. Barreiro, Z. P. Johnson, K. D. Hansen, V. Michopoulos, D. Toufexis, … and Y. Gilad. 2012. Social environment is associated with gene regulatory variation in the rhesus macaque immune system. Proceedings of the National Academy of Sciences 109:6490– 6495.

Tung, J., J. E. Charpentier, D. A. Garfield, J. Altmann, and S. C. Alberts. 2008. Genetic evidence reveals temporal change in hybridization patterns in a wild baboon population. Molecular Ecology 17:1998–2011.

Vilgalys, T. P., A. S. Fogel, J. A. Anderson, R. S. Mututua, J. K. Warutere, I. L. I. Siodi, … and J. Tung. 2022. Selection against admixture and gene regulatory divergence in a long-term primate field study. Science 377:635–641.

Wasser, S. K., G. W. Norton, S. Kleindorfer, and R. J. Rhine. 2004. Population trend alters the effects of maternal dominance rank on lifetime reproductive success in yellow baboons (*Papio cynocephalus*). Behavioral Ecology and Sociobiology 56:338–345.

Weingrill, T. 2000. Infanticide and the value of male-female relationships in mountain chacma baboons. Behaviour 136:337–359.

Wrangham, R. W. 1980. An ecological model of female-bonded primate groups. Behaviour 75:262–300.

Zipple, M. N., J. H. Grady, J. B. Gordon, L. D. Chow, E. A. Archie, J. A. Altmann, and S. C. Alberts. 2017. Conditional fetal and infant killing by male baboons. Proceedings of the Royal Society B 284:20162561.

