## Supplementary Materials for "Activity budgets, social behavior, and fitness outcomes associated with a group fusion in baboons (*Papio cynocephalus* x *P. anubis*)"

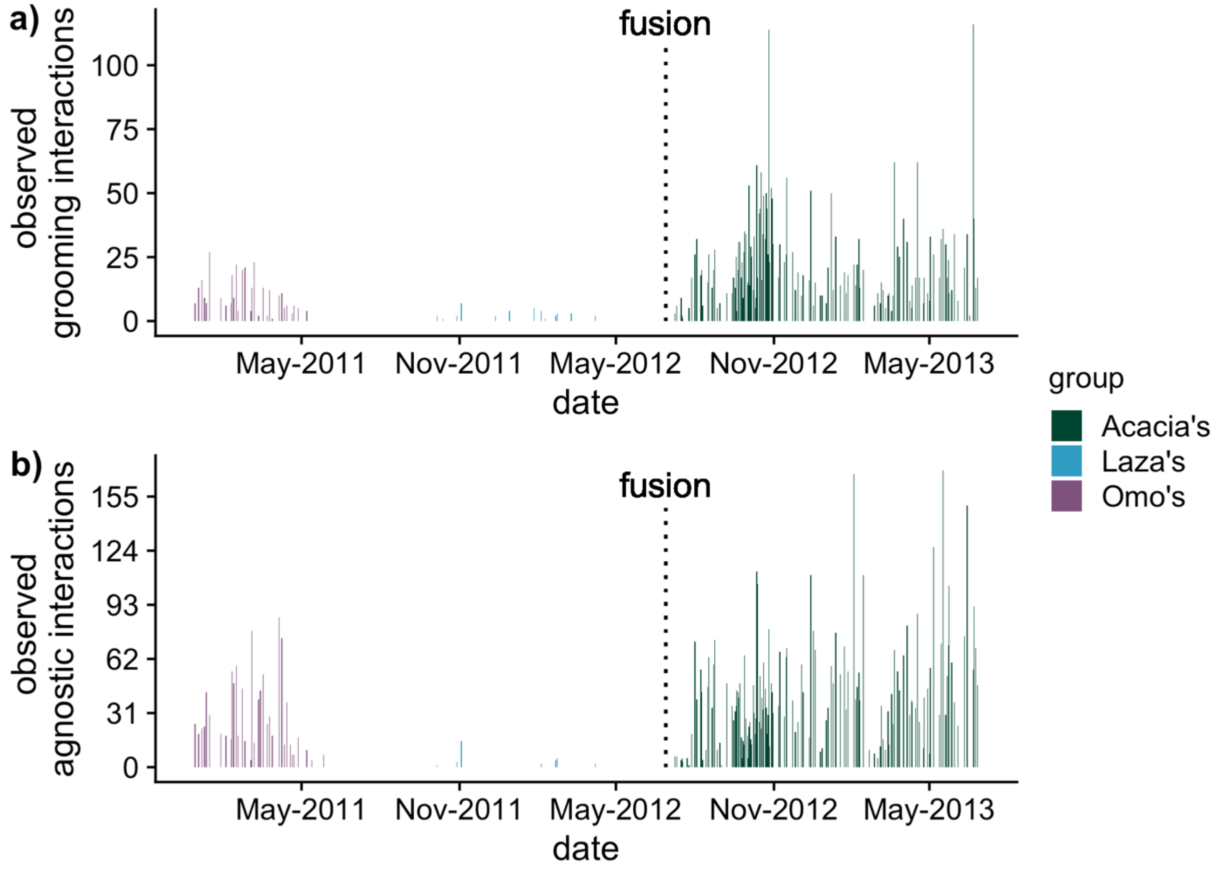


Fig. S1: Availability of social data for baboon groups in the Amboseli basin of Kenya before and after they fused into Acacia's group (data range from December 28th of 2010 to June 26th of 2013; 18 months before the fusion to one year after). Bars show the total number of observed a) grooming interactions between all group members or b) agonistic interactions recorded between all group members in each group per day over time. Behavioral data collection for Omo's group stopped when they were dropped as a study group in May of 2011, although censuses continued to be collected 1-4 times per month until the fusion. Behavioral data for Laza's group were collected infrequently as they were in the process of gradually being dropped as a study group after splitting from a larger group in October of 2011; they were censused ~3 times per week between October 2011 and the fusion.

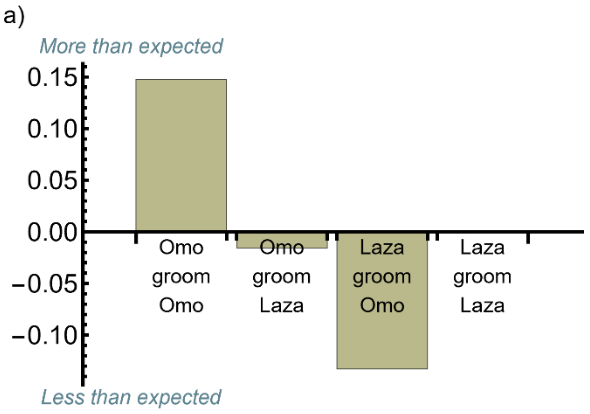


Fig. S2: Distribution of adult female-female a) grooming and b) agonistic interactions in the three-month period between June 29$^{th}$ and September 29$^{th}$ following the fusion of Laza's and Omo's groups in the Amboseli basin of Kenya. The null expectation that interactions between female baboons from each group would be proportional to the number of dyads with a given set of pre-fusion group origins is shown as y=0. Because there were 5 females in Omo's group, 3 in Laza's group, and females cannot interact with themselves, the expected proportions were: Omo-to-Omo: 20/56, Omo-to-Laza: 15/56, Laza-to-Omo: 15/56, and Laza-to-Laza: 6/56. The y-axis values for each bar show the difference between the observed and expected distributions for a) grooming and b) agonistic interactions. For example, a value of 0.2 indicates that a behavior was 20% more likely to be observed than expected based only on the frequency of females from each initial group.

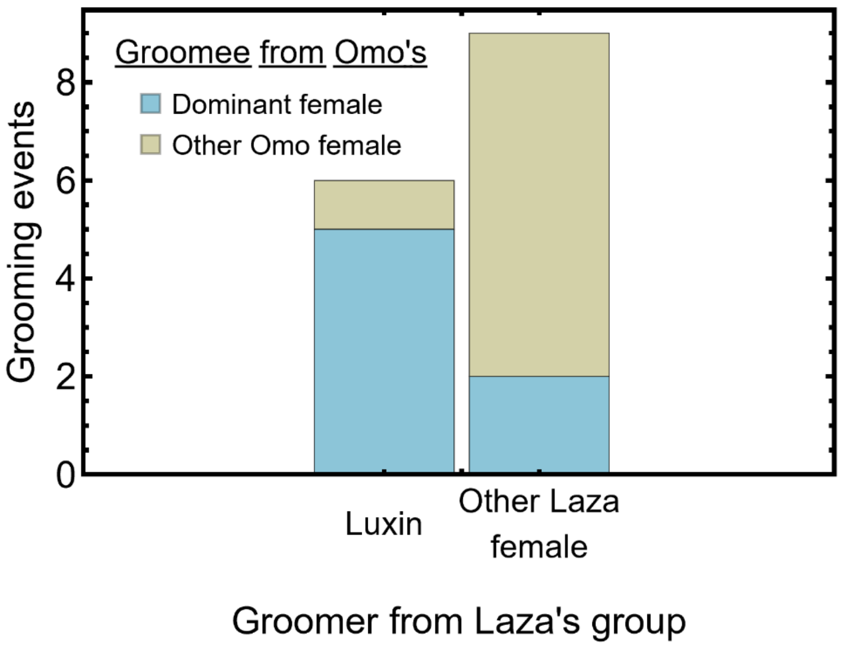


Fig. S3: Number of grooming events initiated by Luxin, a specific female from Laza's group, to female baboons in Omo's group (left bar) versus other females from Laza's group (right bar) in the three months between June 29th and September 29th following the fusion of Laza's and Omo's groups in the Amboseli basin of Kenya. Color indicates the identity of the grooming recipient (blue = the dominant female, Acacia; tan = any other female from Omo's group). Note that more than half of all grooming events by the other two Laza's females come in the last week of September, suggesting that Luxin invested in grooming Omo's females before other females originating from Laza's group did.

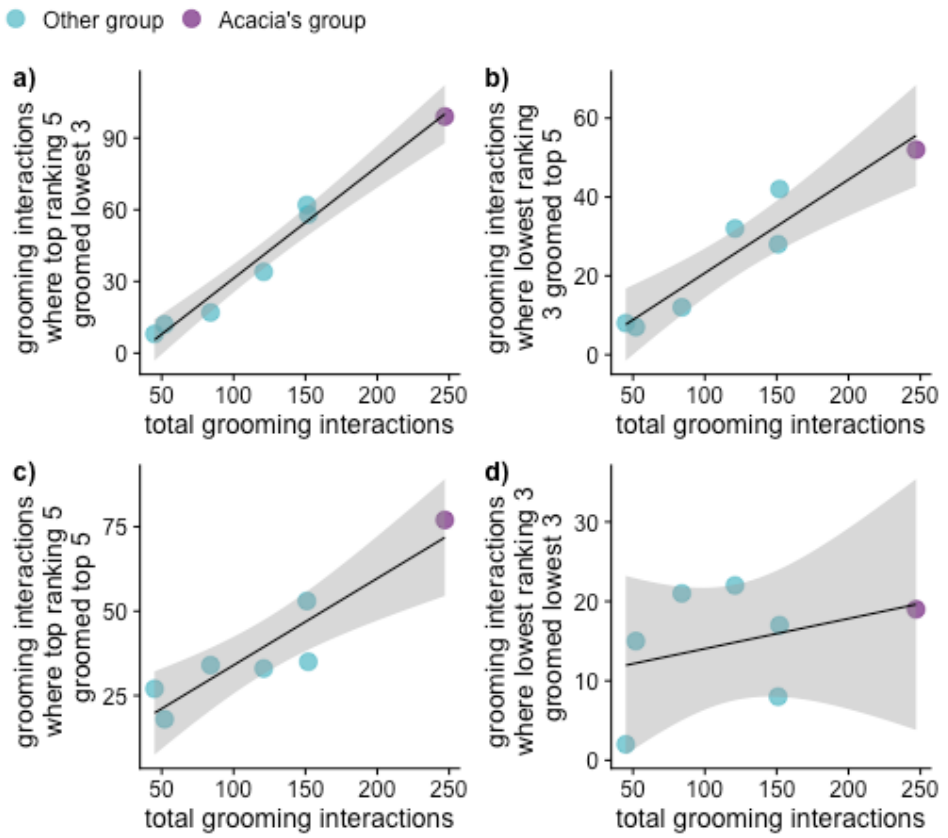


Fig. S4: Relationship between total grooming interactions and grooming interactions between specific subsets of group members, based on rank, in groups containing eight adult female baboons between September 29th and December 29th of a given year (between 1972 and 2023) in the Amboseli basin of Kenya: a) one of the five highest-ranking females groomed one of the three lowest-ranking females, b) one of the three lowest-ranking females groomed one of the five highest-ranking females, c) one of the five highest-ranking females groomed one of the five highest-ranking females, and d) one of the three lowest-ranking females groomed one of the three lowest-ranking females. Acacia's group in the four to six months after it formed via fusion is plotted in purple while other groups are plotted in blue. The solid black line shows the linear regression trendline from a regression where the number of grooming interactions falling within a particular category was regressed on the total number grooming interactions. The shaded gray area shows the 95% confidence interval around the regression line. Note that this figure parallels Fig. 4 but includes data further removed from the fusion. Comparisons are made between the top five and bottom three females because the three lowest ranking adult females in Acacia's group were in Laza's group prior to the fusion and the five highest ranking adult females were in Omo's group prior to the fusion (Fig. 1). Note that the vertical axis scale changes across panels.

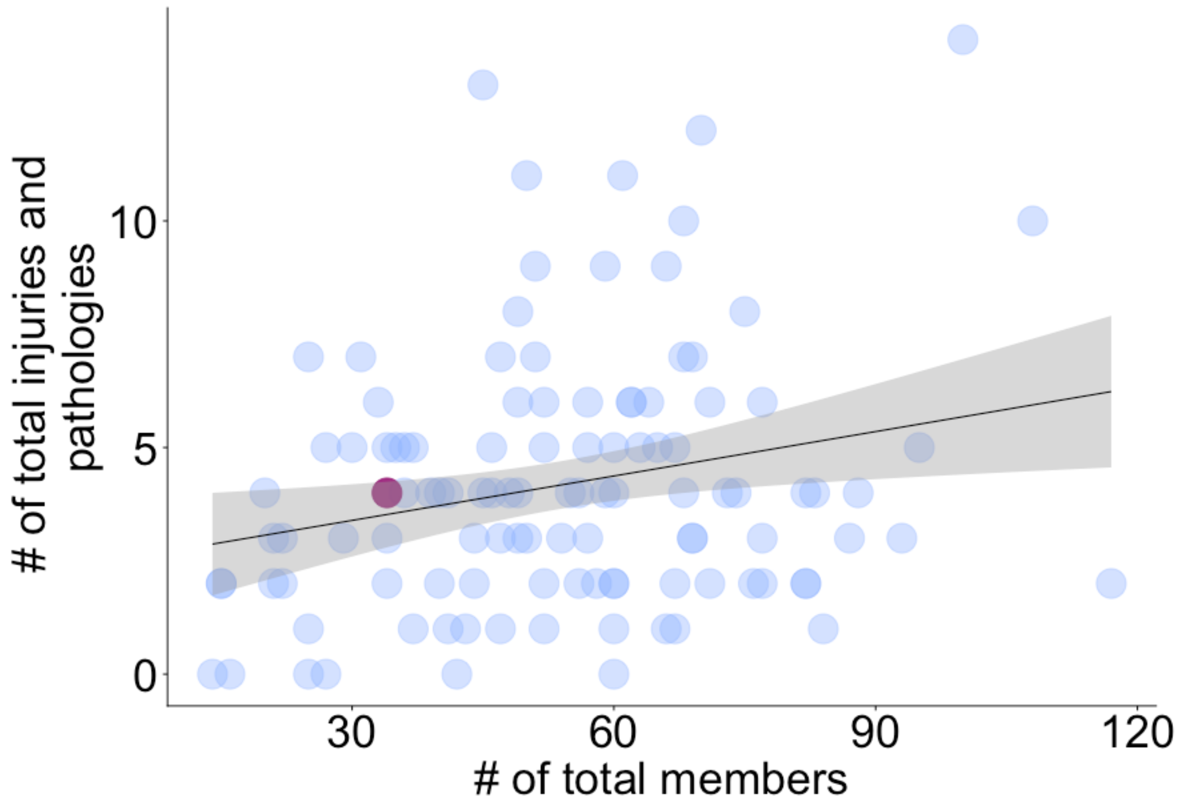


Fig. S5: Number of recorded injuries and pathologies affecting all baboons in a group between June 29th and September 29th of a given year (from 2000 to 2023) in the Amboseli basin of Kenya versus the total number of members in the group (adults of both sexes, juveniles, and infants). Acacia's group in the three months after it formed via fusion is plotted in purple while other group-year combinations are plotted in blue. Note that this figure parallels Fig. 5 but includes data injuries and pathologies for all individuals instead of only adult females.
